# Distinct gut microbiome patterns associate with consensus molecular subtypes of colorectal cancer

**DOI:** 10.1101/153809

**Authors:** Rachel V Purcell, Martina Visnovska, Patrick J Biggs, Sebastian Schmeier, Frank A Frizelle

**Affiliations:** Department of Surgery, University of Otago, Christchurch, New Zealand.; Institute of Natural and Mathematical Sciences, Massey University, Auckland, New Zealand.; Hopkirk Institute, Institute of Veterinary, Animal and Biomedical Sciences, Massey University, Palmerston North, New Zealand.

**Author notes:** **Corresponding author:** Rachel Purcell, PhD Department of Surgery, University of Otago, Christchurch PO Box 4345 Christchurch New Zealand.

**Keywords:** Microbiome, colorectal cancer, molecular classification, oral pathogens, CMS1

## Abstract

Colorectal cancer (CRC) is a heterogeneous disease and recent advances in subtype classification have successfully stratified the disease using molecular profiling. The contribution of bacterial species to CRC development is increasingly acknowledged, and here, we sought to analyse CRC microbiomes and relate them to tumour consensus molecular subtypes (CMS), in order to better understand the relationship between bacterial species and the molecular mechanisms associated with CRC subtypes. We classified 34 tumours into CRC subtypes using RNA-sequencing derived gene expression and determined relative abundances of bacterial taxonomic groups using 16S rRNA amplicon metabarcoding. 16S rRNA analysis showed enrichment of Fusobacteria and Bacteroidetes, and decreased levels of Firmicutes and Proteobacteria in CMS1. A more detailed analysis of bacterial taxa using non-human RNA-sequencing reads uncovered distinct bacterial communities associated with each molecular subtype. The most highly enriched species associated with CMS1 included *Fusobacterium hwasookii* and *Porphyromonas gingivalis*. CMS2 was enriched for *Selenomas* and *Prevotella* species, while CMS3 had few significant associations. Targeted quantitative PCR validated these findings and also showed an enrichment of *Fusobacterium nucleatum, Parvimonas micra* and *Peptostreptococcus stomatis* in CMS1. In this study, we have successfully associated individual bacterial species to CRC subtypes for the first time.

## Introduction

Colorectal cancer (CRC) is one of the most common cancers, as well as one of the cancers with the highest mortality worldwide[1]. CRC is a highly heterogeneous disease, with varying clinical outcomes, response to therapy, and morphological features. Therefore, classification into clinically useful and reproducible subtypes has been a goal within the research community for many years. Studies of molecular features, such as *BRAF*, *KRAS and TP53* mutation status, microsatellite instability (MSI), CpG island methylator phenotype (CIMP), somatic copy number alterations (SCNA), and activation of various molecular pathways such as WNT and MYC, have been used with some success to stratify CRC into subgroups[2-5]. The advent of large-scale sequencing technologies has recently facilitated the development of a Consensus Molecular Subtyping (CMS) system for CRC based solely on tumour gene expression[6]. The strong association of these CMS subtypes with distinct molecular features and pathway activation provides an indication of potential mechanisms underlying the disease.

Most CRCs are sporadic and follow a pattern one would expect from a yet unidentified environmental source. The human colon plays host to a vast and complex microbial community of < 10^12^ microorganisms[7], and a growing body of evidence points to a role for gut microbial dysbiosis in the development of CRC[8]. Comparison of faecal microbiomes from CRC patients and healthy controls[9-12] has identified particular bacterial species that are enriched in CRC, and analysis of tumour, adenoma, and matched normal tissue from the same patients found that changes in local communities of potentially interacting bacterial taxa are associated with different disease states[12, 13]. Correlations between particular cancer mutations and changes in microbial communities, and transcriptional remodelling associated with specific bacteria have recently been described in CRC[14, 15]. However, a global investigation of the association of tumour gene expression with tumour metagenomics has yet to be described. Identification of specific species or bacterial communities associated with CRC subtypes could facilitate improved screening and diagnostics, and understanding the underlying pathogenic mechanisms will pave the way for the development of targeted interventions, such as microbiome modulation and vaccines for CRC prevention.

Here, we looked at gene expression and related CMS subtypes of a cohort of 34 CRC patients using data derived through RNA-sequencing, and use both bacterial 16S rRNA gene analysis and interrogation of non-human RNA sequences from tumour tissue to establish metagenomic profiles for each tumour sample. The combination of tumour transcriptomics and metagenomics has allowed us to identify bacterial species with potential roles in the mechanisms underlying particular molecular subtypes of CRC.

## Materials and Methods

### Patient cohort and samples

Colorectal cancer tumour samples were collected from 34 patients during surgical resection of previously untreated tumours. Samples were collected with patients' written, informed consent, and this study was carried out with approval from the University of Otago Human Ethics Committee (ethics approval number: H16/037). Twenty of the patients were female, and patient ages at the time of surgery ranged from 44–88 years (mean age, 74 years, see Table 1). One tumour sample was from the rectum, 21 tumours were from the right side of the colon, and 12 from the left side. One sample was of a large colorectal adenoma, not an adenocarcinoma. Histologically, four tumours were described as well differentiated, 20 were moderately and nine were poorly differentiated. Two tumours showed signet-ring histology and three were mucinous type. Postoperative staging showed that five tumours were stage 1, 14 and 13 tumours were stage 2 and 3, respectively, while only one tumour was stage 4. Patient characteristics are given in more detail in Table 1.

**Table 1.** Patient cohort characteristics. *post-operative; ** including one large adenoma; n, number of patients

|  | Patients (n) |
| --- | --- |
| **Age** |  |
| 44-88 years |  |
| Mean = 74 years |  |
| **Gender** |  |
| Males | 14 |
| Females | 20 |
| **Site** |  |
| Right | 21** |
| Left | 12 |
| Rectum | 1 |
| **Stage\*** |  |
| 1 | 5 |
| 2 | 14 |
| 3 | 13 |
| 4 | 1 |
| **Differentiation** |  |
| Well | 4 |
| Moderate | 20 |
| Poor | 9 |
| **Histology** |  |
| Signet-ring cell | 2 |
| Mucinous | 3 |
| Lymphovascular invasion | 16 |
| Extramural venous invasion | 7 |
| Perineural invasion | 5 |
| Lymph node positivity | 13 |

### Nucleic acid extraction

Samples were immediately frozen in liquid nitrogen and initially stored at -80°C. They were subsequently transferred to RNAlater ICE™ (Qiagen), and stored at - 20°C, prior to nucleic acid extraction. RNA was extracted from < 20 mg of tissue using RNEasy Plus Mini Kit (Qiagen), including DNAse treatment, following tissue disruption using a Retsch Mixer Mill. DNA was extracted using DNeasy Blood and Tissue Mini Kit (Qiagen), also following tissue disruption. DNA extraction included overnight incubation with proteinase K, and treatment with RNAse A. Purified nucleic acids were quantified using the NanoDrop 2000c spectrophotometer (Thermo Scientific, Asheville, NC, USA), and stored at -80°C. Nucleic acids were extracted from all tumour samples in a single batch by one operator, to avoid inter-batch variation.

### RNA sequencing

Sample preparation, including library creation and ribosomal RNA depletion (with RiboZero Gold) was carried out using Illumina TruSeq V2 reagents. RNA-sequencing was carried out using the Illumina HiSeq 2500 V4 platform to produce 125bp paired end reads. The libraries were sequenced on two lanes of the HiSeq instrument. To avoid technical biases caused by sequencing the libraries on different lanes, each sample library was split equally to the two lanes. Sequences from the two lanes were merged for each sample during the data processing phase.

### Quality control and gene expression quantification

In order to calculate a gene expression profile for each sequenced sample and later on to classify the samples into CMS subtypes, the raw sequenced reads were processed in the following way: First, low quality segments of reads as well as remnants of adapter sequences were removed and very short reads were discarded. Second, the reads passing the previous step were mapped to a reference human genome sequence and a read count for every annotated gene was calculated for each sample. The read counts were later transformed to gene expression profile expressed by a measure 'transcripts-per-million' (TPM). Last, the CMS classifier[6] was used to assign a molecular subtype of the disease to each sample based on the gene expression profiles.

Adapter sequences were removed using utility fastq-mcf (v1.1.2.537) from EA Utils[16]. Next, SolexaQA++ (v3.1.6)[17] was used to trim low-quality segments of the reads and only sufficiently long reads were kept for further analysis. At the first step SolexaQA++ dynamictrim stored for each read only the longest continuous segment, such that the probability of a base being called in error was less than 0.01. Afterwards, segments shorter than 50bp were discarded using SolexaQA++ lengthsort.

The human genome reference sequence GRCh38 and HAVANA annotation were used as the reference genome to map the cleaned reads to using the STAR (v2.5.2b) mapping tool[18]. After the mapping stage, reads from two sequencing lanes were merged together using samtools merge (v1.3.1)[19]. Later, a read count table was created using htseq-count (v0.6.1p1)[20] for each sample and the raw read counts were transformed to TPM values using R and Bioconductor package DESeq2 (v1.10.1)[21].

### CMS classification

The CRC Subtyping Consortium created the consensus molecular subtype (CMS) classification by utilizing six previously established classification systems. Patients' gene expression profiles (n = 3962) were collected from various public and proprietary datasets. The profiles were classified due to each of the previously known systems and a graph of relationships among subtypes of the various classification systems was used in a Markov Cluster Algorithm to identify a number of consensus subtypes. A smaller set of core consensus patient data (n = 3104) and their gene expression profiles were used to train the Random Forest based CMS classifier.

Besides the random forest classifier, the authors made a Single Sample Predictor (SSP) method available in the CMS classifier (v1.0.0, https://www.synapse.org/#!Synapse:syn4961785) R package, and this method was used to classify our samples into molecular subtypes of colorectal cancer. The SPP method is less sensitive to differences in normalisation techniques used during the data processing and in terms of its performance comparable to the random forest classifier.

In order to classify a sample to one of the four subtypes, the SSP method calculates similarity of the sample to a centroid of each subtype. If the sample shows acceptable level of similarity to exactly one of the centroids, the subtype corresponding to the centroid is assigned to the sample undergoing the classification. Similarity between two gene expression profiles is calculated as Pearson's correlation of log2 scaled values from the profiles. A sample is considered to be similar to a centroid if the correlation is at least 0.15. In order for a sample to be classified, a correlation to the most similar centroid has to be higher by 0.06 than a correlation to the second most similar centroid. These values are set by default in the SSP method. Centroid genome expression profiles are hard-coded in the method as information from the random forest classifier and the training data were used to calculate the values.

### Metabarcoding by 16S rRNA

Libraries containing16S rRNA were prepared with 20 ng of DNA for each sample using primer pairs flanking the V3 and V4 regions of the 16S rRNA gene (16SF_V3: 5′-TATGGTAATTGGCCTACGGGAGGCAGCAG-3' and 16SR_V4: 5′-AGTCAGTCAGCCGGACTACHVGGGTWTCTAAT-3′), and Illumina sequencing adaptors and barcodes added using limited cycle PCR (40 cycles). Amplicon sequencing was carried out using the Illumina MiSeq platform, and paired end reads of length 250bp were generated.

During data processing, short overlapping forward and reverse reads coming from the same fragment were joined together by FLASh (v1.2.11)[22] to form overlapped sequences of the V3-V4 16S region. After joining, the resulting fragments were trimmed that the probability of a base being called in error was less than 0.01, and minimal length of a fragment was at least 50bp. This step was done by SolexaQA++ (v3.1.5). Next, chimeric sequences were removed by the QIIME (v1.9)[23] scripts identify_chimeric_seqs.py and filter_fasta.py. A collection of sequences suitable for further QIIME analysis was thus obtained. Later on, the script pick_de_novo_otus.py was used to identify *de novo* operational taxonomic units (OTUs) and to link the OTUs to the available bacterial taxonomy. Taxa were then summarized for various metadata classes (e.g. CMS, tumour location, etc.) using summarize_taxa_through_plots.py.

### Taxonomic classification of RNA-sequencing reads using Kraken

All NCBI Refseq bacterial genomes with "Complete Genomes"- or "Chromosome"- level genomes were downloaded from NCBI FTP site (ftp://ftp.ncbi.nlm.nih.gov/genomes/refseq/bacteria/) based on information in the "assembly_summary.txt" file as of 19th January 2017. A list of the genomes can be accessed in Supplementary Table S1. Additional genomes known to play role in CRC were added disregarding their genome status (see Supplementary Table S2). Using the genome fasta-files, a new Kraken database (https://ccb.jhu.edu/software/kraken/) was created using "kraken-build --build" with default parameters[24]. The resulting Kraken database had a size of ~131GB. All RNA-seq reads that were unable to be mapped to the human reference genome (GRCh38) were extracted per sample and were used as input to Kraken (v0.10.6) using our custom Kraken database for taxonomic classification. This resulted in bacterial abundances per CRC sample based on unmapped RNA-seq reads. We visualized bacterial abundances per CRC subtype, by combining all reads of all samples of a CRC subtype and those reads were input to Krona (v2.7). Interactive plots are available at https://crc.sschmeier.com.

### Differential bacterial abundances in CMS subtypes

To identify bacterial strains that are enriched or depleted in one CMS subtype compared to all other subtypes, we employed a strategy similar to common differential expression analyses. We term this approach "differentially expressed taxa". Using edgeR Bioconductor package (v3.14.0) [25], we identified bacterial taxa whose abundances are significantly different among CMS subtypes.

In this analysis, for each CRC sample we used the assigned CMS subtype together with a list of bacterial taxa identified by Kraken and corresponding raw read counts as input data. We treated all samples of a certain CMS subtype as replicates belonging to the subtype. We ran differential analysis of each CMS subtype against all the other classified samples (more information can be found at https://gitlab.com/s-schmeier/crc-study-2017). This resulted in bacterial taxa that are enriched (or depleted) in a subtype as compared to all other subtypes. We performed this analysis on the species-level taxa for all CMS subtypes.

### Quantitative PCR

Levels of the *Porphyromonas gingivalis, Bacillus coagulans, Selenomas sp., F. nucleatum*, *P. stomatis*, *P. micra* and the reference gene, prostaglandin transporter (*PGT*)[26] were simultaneously measured from genomic DNA extracted from CRC tumour samples, using qPCR on a LightCycler^®^480 thermocycler (Roche Diagnostics, Indianapolis, IN, USA), as previously described[27]. Primers for each gene are given in Supplementary Table S3. Genomic DNA from purified bacterial samples were used as positive controls (DSMZ, Germany). The levels of each of the bacterial species in each DNA sample were calculated as a relative quantification (RQ). Calculations were made using 2^-ΔCt^, where ΔCt is the difference in Ct values between the gene target of interest and reference gene for a given sample.

### Differential gene expression analysis

We used count tables from the RNA sequencing mapping and counting procedure (see above), in addition to the CMS subtype information produced through the CMS classification step, and ran a differential expression analysis for all genes. We used each sample within a given subtype as a replicate of that subtype, and ran edgeR (v3.14.0) to compare each subtype against all other samples not in the CMS subtype, e.g. samples in CMS1 against all other classified samples (samples in CMS2 plus CMS3). We extracted genes that are up- or down-regulated, using a Benjamini and Hochberg false-discovery rate (FDR)[28] adjusted *P*-value (< 0.1), and a log2 fold-change greater or smaller than zero (more information can be found at https://gitlab.com/s-schmeier/crc-study-2017).

### Gene-set enrichment analysis

We used gene-sets per CMS subtype, and the type of expression (i.e. up- or down-regulation) as input, and calculated a *P*-value for enrichment of the gene-set in biological categories, using Fisher's exact test for count data ("fisher.test" method in R). All *P*-values were adjusted for multiple testing using the false-discovery adjustment method from Benjamini & Hochberg, using R-method "p.adjust" (more information can be found at https://gitlab.com/s-schmeier/crc-study-2017). The biological categories and corresponding gene-sets used in the analysis were extracted from MSigDB[29] (version 5.2). We sub-selected the following categories for the analysis: KEGG, REACTOME, BIOCARTA, PID, HALLMARK GENES, and Gene Ontology (GO) biological processes. The background set of genes for each gene-set enrichment analysis (GSEA) test was all genes associated with any of the above categories.

## Results

### Classification of CRC samples into consensus molecular subtypes

Reads generated by RNA-sequencing were quality checked, mapped to the human reference genome, and gene expression was quantified based on the number of reads mapped to particular transcript models. The RNA from one tumour sample was too degraded to carry out RNA-sequencing, leaving 33 samples. Gene expression profiles from each patient were used as input data to the publicly available CMS subtype classifier[6] (see Materials and Methods). The CMS subtype for each sample based on the classification was recorded. Five CRC samples were designated as unclassified. The proportion of CRC samples in each subgroup is shown in Table 2, compared with that seen in the CMS classification by the CRC subtyping consortium[6].

**Table 2.** Comparison of proportion of patients in each consensus molecular subtype (CMS) and unclassified (UC) tumours from this study and the original classification study by the CRC subtyping consortium.

|  | **CMS1** | **CMS2** | **CMS3** | **CMS4** | **UC** |
| --- | --- | --- | --- | --- | --- |
| **This study** | 18% | 39% | 27% | 0% | 15% |
| **CMS study** | 14% | 37% | 13% | 23% | 13% |

### Biological features of CMS subtypes

In order to validate the CMS classification approach, we identified differentially expressed genes (DEGs) for groups of samples belonging to each subtype (CMS) by comparing the samples of a subtype against all other samples not belonging to that subtype. We did this for all subtypes in which we classified our samples (CMS1, CMS2, and CMS3). We split the resulting DEG into up- and down-regulated genes and ran gene-set enrichment analysis (GSEA) with each individual set of genes to identify categories in which these genes are enriched (see Materials and Methods). Although we were not able to exactly reproduce the methodology used in the publication describing the original classification[6], the enriched biological categories per CRC subtype in our data closely follow the originally identified categories, confirming that our approach for classifying the CRC samples was successful (see Supplementary Table S4).

Strikingly, most highly ranked categories for up-regulated DEGs of CMS1 show clear immunological signatures, e.g. immune response, interferon-γ response, inflammatory response, TNFα signalling through NF_K_β and cytokine-mediated signalling (see Supplementary Table S4). Downregulated DEGs in CMS1 are involved in pathways associated with digestion, metabolism and the negative regulation of morphogenesis.

Up-regulated DEGs of CMS2 show significant enrichment in cell-cycle signatures, e.g. DNA damage checkpoint, DNA replication and synthesis and cell cycle regulation. DEGs associated with Myc signalling were also enriched, although statistical significance was not reached. However, no enrichment of Wnt-associated signalling was found in CMS2 in our cohort. Interestingly, the down-regulated DEG in CMS2 show a greater number of significantly enriched categories, especially those associated with immunological signatures, which agrees with the observation made in the study by Guinney et al[6].

GSEA of our designated CMS3 samples showed many highly significant categories associated with metabolism, e.g. lipid and steroid metabolism, amongst others. Interestingly, we found that many down-regulated DEGs are enriched in cell-cycle related categories, and those involved in DNA replication, synthesis and damage checkpoints.

Despite differences in the methodology of the analysis, as well as in the biological categories used, overall, we recovered many similar biological signatures through GSEA of our DEG groups per subtype, suggesting that the originally proposed classification system is appropriate, and our analysis produced comparable results.

### Insights into 16S rRNA metabarcoding data

Metabarcoding analysis of microbial communities localized on tumour tissue samples based on the V3-V4 hypervariable region of the 16S rRNA gene was carried out using QIIME. Representative sequences were identified for each operational taxonomic unit (OTU), and these sequences were used to assign taxonomy to each OTU using the Greengenes reference OTU database. Relative abundances of taxonomic groups in each CRC sample was calculated on various taxonomic levels (from phyla to genera), and the bacterial phyla present on average with abundance over 1% are listed for each sample from our cohort in Table 3. It can be seen that the samples differ remarkably already in abundances at the level of bacterial phyla. In Figure 1, we show the relative abundances for various groups of samples from our cohort. As for CMS subtypes, visible differences are in enrichment of Fusobacteria and Bacteroidetes, and decreased levels of Firmicutes and Proteobacteria in CMS1, compared to CMS2 and CMS3 (Figure 1A).

Analysis of differences in taxonomic abundances between samples based on other clinicopathological variants, such as tumour differentiation (Figure 1B) and tumour location (Figure 1C) also showed changes at the phylum level; decreased levels of Fusobacteria and Firmicutes, and increased abundance of Proteobacteria and Bacteroides were seen with decreasing tumour differentiation, while right-sided tumours had increased Fusobacteria compared to left-sided tumours.

**Figure 1.**
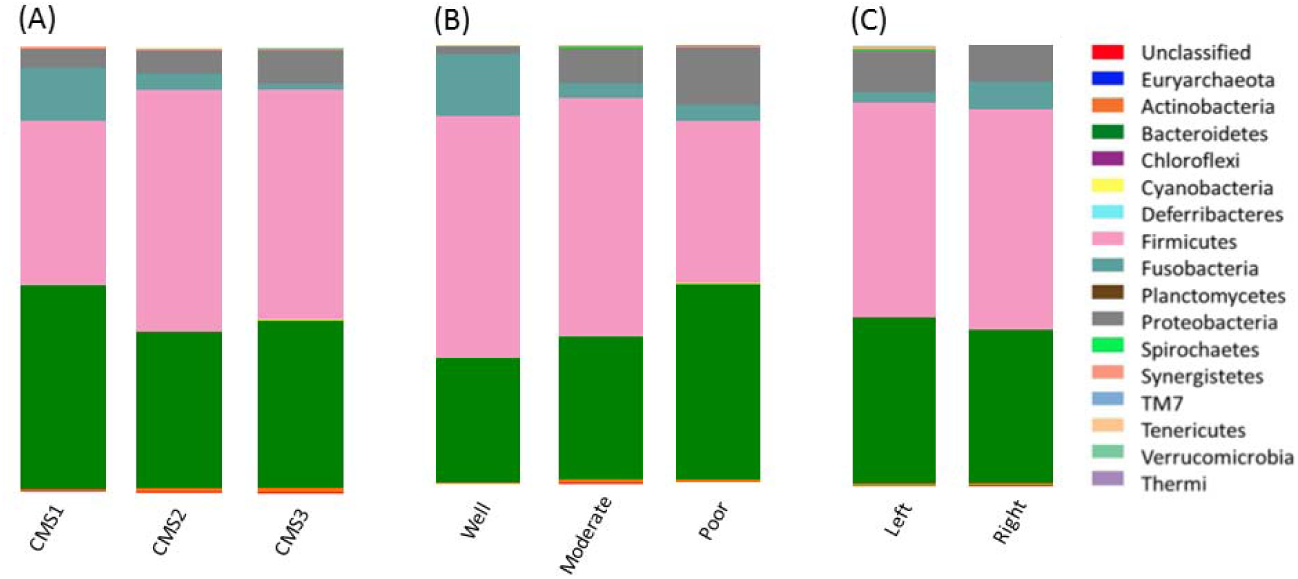
Phylum-level bar chart grouped by (A) consensus molecular subtype (CMS) group, (B) histological tumour differentiation and (C) location of tumour. Bar charts are colour coded by phyla.

**Table 3.** Phyla of bacteria present in each sample, with abundance > 1%, as per 16S rRNA analysis. SD; standard deviation.

| Sample | Firmicutes | Bacteroidetes | Proteobacteria | Fusobacteria |
| --- | --- | --- | --- | --- |
| CRC1 | 49.18 | 41.13 | 4.56 | 1.67 |
| CRC2 | 29.84 | 41.03 | 5.27 | 17.98 |
| CRC3 | 50.12 | 1.05 | 46.57 | 0.01 |
| CRC4 | 60.56 | 25.04 | 12.87 | 0.79 |
| CRC5 | 62.8 | 28.87 | 7.34 | 0.27 |
| CRC6 | 27.31 | 56.49 | 15 | 0.02 |
| CRC7 | 31.98 | 60.06 | 5.39 | 2.11 |
| CRC8 | 69.09 | 27.25 | 2.49 | 0 |
| CRC9 | 15.3 | 60.37 | 2.7 | 20.95 |
| CRC10 | 61.35 | 24.89 | 10.58 | 1.41 |
| CRC11 | 35.13 | 51.33 | 6.48 | 0 |
| CRC12 | 58.67 | 38.42 | 1.3 | 1.13 |
| CRC13 | 76.32 | 18.58 | 2.12 | 2.01 |
| CRC14 | 37.57 | 20.7 | 39.93 | 1.11 |
| CRC15 | 54.01 | 38.21 | 4.93 | 0.71 |
| CRC16 | 34.76 | 33.92 | 19 | 11.98 |
| CRC17 | 64.94 | 29.15 | 5.44 | 0 |
| CRC18 | 72.84 | 20.91 | 4.48 | 0 |
| CRC19 | 69.78 | 19.28 | 6.69 | 3.87 |
| CRC20 | 52.87 | 35.38 | 10.35 | 0 |
| CRC21 | 42.27 | 49.98 | 5.64 | 0.93 |
| CRC22 | 55.03 | 38.91 | 6 | 0 |
| CRC23 | 56.36 | 33.36 | 7.55 | 1.29 |
| CRC24 | 66.36 | 27.06 | 5.04 | 0.22 |
| CRC25 | 53.77 | 12.03 | 0.34 | 33.65 |
| CRC26 | 71.38 | 26.3 | 0.45 | 0.17 |
| CRC27 | 11.06 | 84.94 | 0.91 | 3.09 |
| CRC28 | 58.42 | 28.99 | 10.37 | 0 |
| CRC29 | 35.78 | 61.23 | 1.93 | 0.84 |
| CRC30 | 36.48 | 59.52 | 1.08 | 1.05 |
| CRC31 | 66.55 | 6.46 | 2 | 23.92 |
| CRC32 | 63.53 | 22.22 | 9.24 | 0.01 |
| CRC33 | 30.18 | 51.72 | 11.12 | 6.66 |
| CRC34 | 15.86 | 80.23 | 0.85 | 0.36 |
| Average (%) | 49.34 | 36.91 | 8.12 | 4.07 |
| SD | 17.92 | 19.39 | 9.97 | 8.05 |

### Taxonomic investigation of non-human RNA-seq reads per CMS subtype

Although 16S rRNA analysis allowed us to identify changes in bacterial communities at the phylum level between CRC subtypes, it did not give us resolution to the species level. In order to identify bacteria that may be associated with particular molecular subtypes at the species level, we performed a taxonomic investigation of non-human RNA-seq reads using Kraken[24] (see Materials and Methods). To validate the consistency of the two approaches, we compared phylum and genus level abundances derived through 16S rRNA analysis or separately derived through non-human mapped RNA-seq reads using Kraken[24]. When correlating the phylum-level abundances derived through 16S rRNA metabarcoding with the abundances derived through Kraken for each CRC sample, we see a high correlation of 85% (Figure 2B). At the genus level, there was an overall lower, but still high correlation as compared to phylum level, showing a reasonable agreement between the 16S rRNA and the RNA-seq Kraken approach (see Figure 2A). However, it should be noted that the number of genera that appear in both 16S rRNA and Kraken-derived data is quite low (76 genera, and only 13 on the phylum level); it is possible that using RNA-seq derived bacterial abundances is more sensitive in detecting taxa than 16S rRNA sequencing, as has been shown previously for whole shotgun sequencing compared to 16S rRNA sequencing[30].

**Figure 2.**
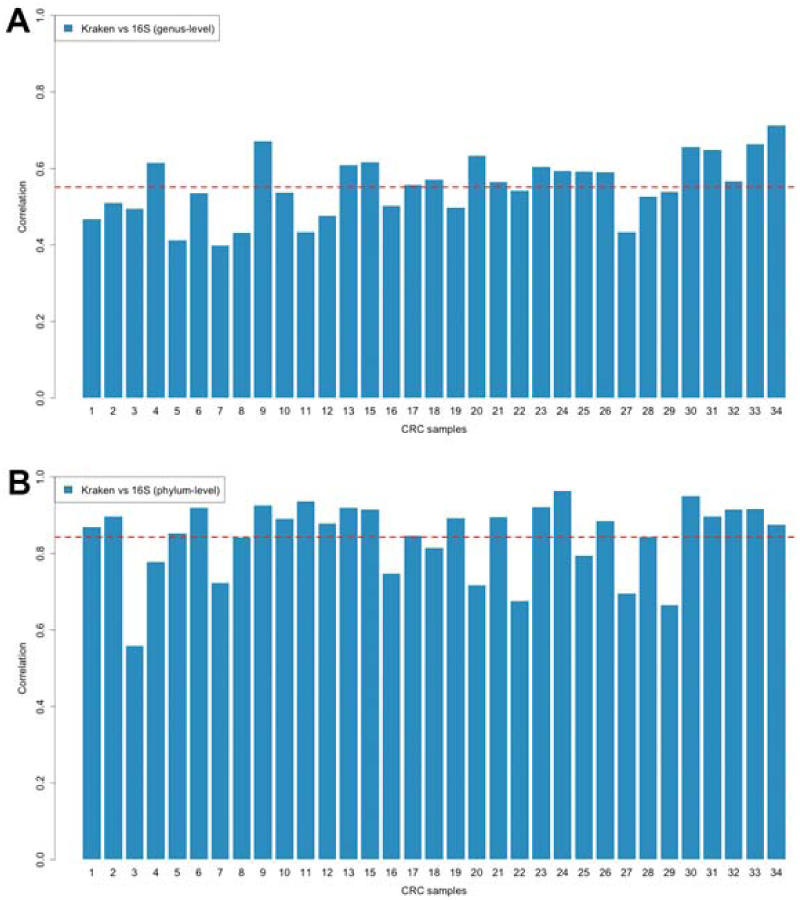
**A**: Correlation between CRC samples using genus-level abundances derived through 16S rRNA metabarcoding and Kraken. 76 genera that appear in both methods were used. **B**: Correlation between CRC samples using phylum-level abundances derived through 16S rRNA metabarcoding and Kraken. Thirteen phyla that appear in both methods were used. Dashed lines indicate the average correlation over all samples.

Analysis of bacterial taxa for each molecular subtype uncovered distinct bacterial communities associated with each CMS subtype (Figure 3 and also https://crc.sschmeier.com for interactive Krona plots). Table 4 shows the 15 most highly enriched genera for each CMS subtype (for a complete list of genera see Supplementary Table S5).

**Figure 3.**
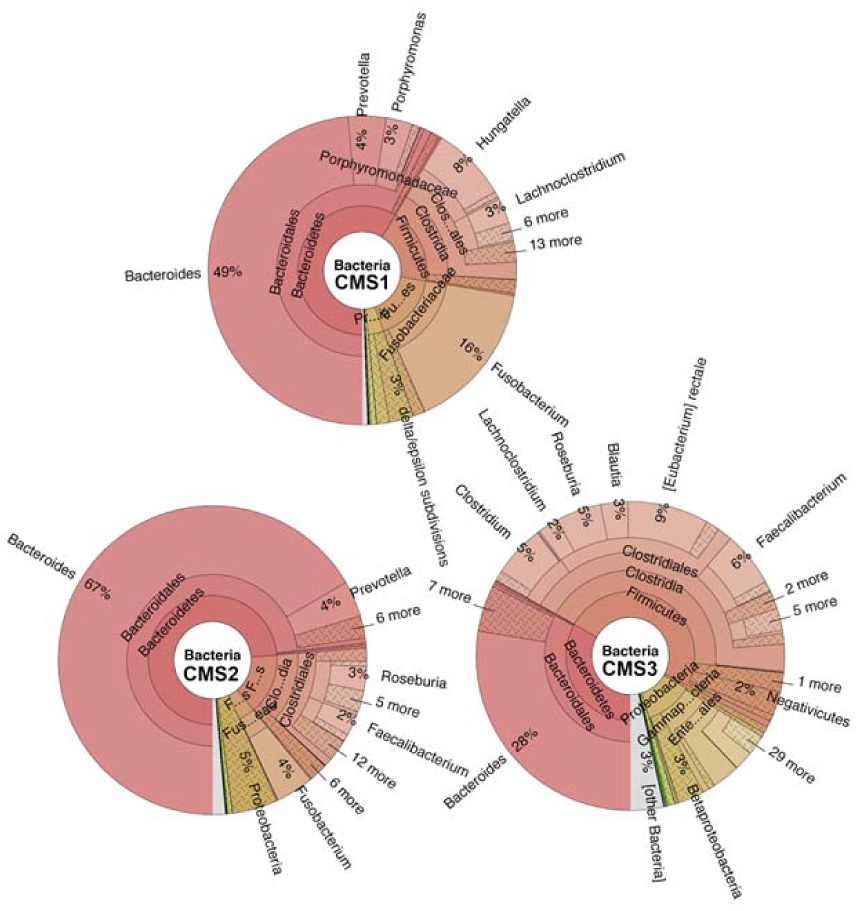
Krona plots of for each CMS showing relative abundance of bacterial taxa at the genus level. Interactive versions of these Krona plots can be further interrogated at https://crc.sschmeier.com.

**Table 4.**
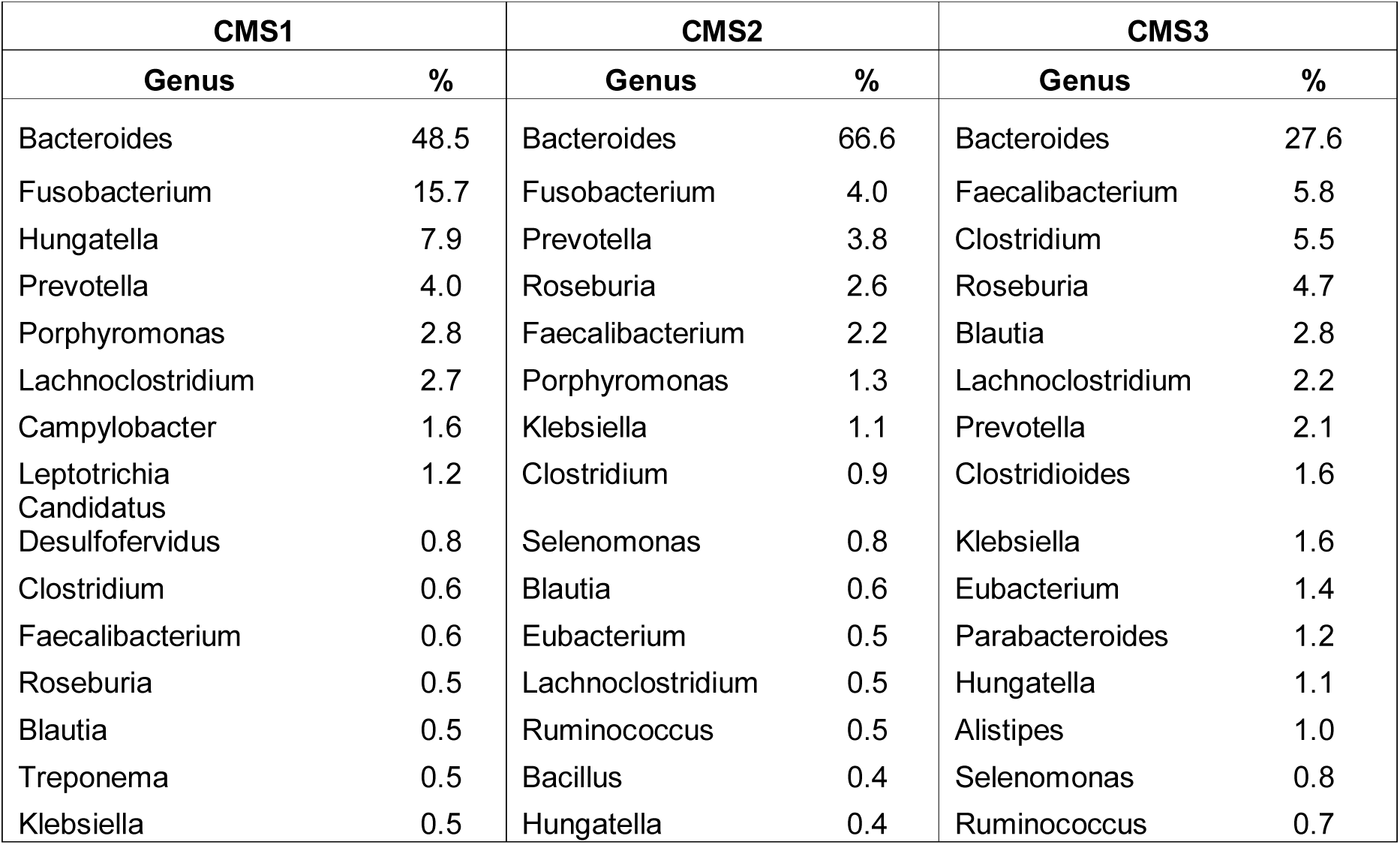
The 15 most highly abundant bacterial genera, as a percentage of the total bacterial genera, for each consensus molecular subtype (CMS), as calculated using RNA-seq metagenomics.

Analysis of Kraken-derived data at the species level revealed that among the highest enriched bacterial species associated with CMS1 were *Fusobacterium hwasooki* (previously *F. nucleatum*) and *Porphyromonas gingivalis*, both known oral pathogens with putative roles in CRC development. *P. gingivalis* is also known to synergistically promote extra-gastrointestinal infections through co-occurrence with *Treponema denticola* and *Tannerella forsythia*, both of which are also strongly associated with CMS1. CMS2 had highly enriched *Selenomonas* and *Prevotella* species, while there were only a few bacterial species significantly associated with CMS3. Of particular interest to this study was the strong association of oral pathogens and bacteria capable of forming biofilms with CMS1. Taken together, the strong immunological and inflammatory signatures associated with this CRC subtype, and the known mechanisms of actions of some of these bacteria in extra-gastrointestinal infections, provides a plausible link between the co-occurrence of certain oral bacteria and the development of CMS1 type CRC.

### Validation of bacterial species in CMS using qPCR

Three bacterial species were chosen for qPCR validation of CMS subtypes using genomic DNA samples extracted from the CRC tumours. The species were chosen based on their high enrichment in a CMS and corresponding low association with the other CMS subtypes. Primer design and commercial availability of bacterial genomic DNA for use as positive controls were also factors in choosing validation targets. *Porphyromonas gingivalis*, *Selenomonas sp*. and *Bacillus coagulans* were chosen due to their strong associations with CMS 1, 2 and 3, respectively. Three further bacteria were targeted due to their known or putative roles in CRC development: the oral pathogens, *F*. *nucleatum*, *Parvimonas micra* and *Peptostreptococcus stomatis*. Relative levels of each species was calculated using ΔCt method, with *PGT* as a reference gene (See Material and Methods). The relative expression values are given in Supplementary Table S6. Relative expression levels were calculated for each molecular subtype compared to the other subtypes and fold-change values generated based on average expression in samples of a subtype to the average expression in all other samples (Figure 4). High abundances of *P. gingivalis* and *Selenomonas sp*. were strongly correlated with CMS1 and CMS2, respectively, validating our findings generated from the Kraken analysis for these two subtypes. The expression of *B. coagulans* was less significantly associated with CMS3 than findings from Kraken analysis suggested. In addition, increased abundances of the oral pathogens, *F*. *nucleatum*, *P*. *micra* and *P*. *stomatis* were associated with CMS1.

**Figure 4.**
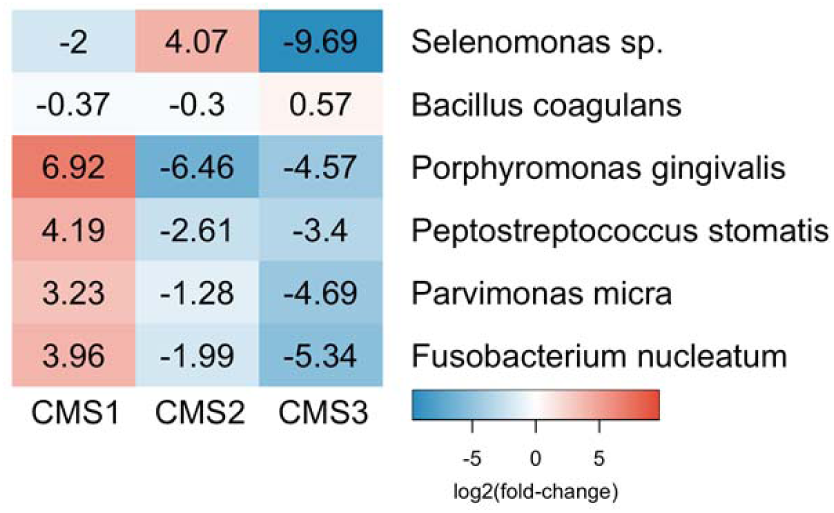
Heatmap of log2 fold-changes in abundance of bacterial targets analysed using qPCR, for each consensus molecular subtype (CMS).

## Discussion

The two main objectives of our study were the validation of molecular subtypes of CRC using published classifiers in an independent CRC cohort, and particularly the examination of differences in bacterial communities associated with different molecular subtypes of CRC. For the first aim, our study found similarities between our cohort and the original classification study of Guinney et al[6] that classified CRCs into consensus molecular subtypes. We found similar proportions of tumours classified as CMS1 and CMS2, in addition to unclassifiable tumours. However, none of the tumours in our study were classified as CMS4, and there was an increased proportion of CMS3 in our cohort compared to that used in the study of Guinney et al: 30% compared to 13%. Several reasons might contribute to the observed differences. First, the small number of patients (33) in our study may have resulted in the absence of CMS4 patients from our cohort; the CRC subtyping consortium used 3104 patients’ samples in their study. Furthermore, differences in analytical methods used on RNA-sequencing data might be a contributing factor. We were unable to exactly replicate the methodology of the original CRC subtyping consortium study for several reasons. First, the original study made use of data from The Cancer Genome Atlas (TCGA, http://cancergenome.nih.gov/), which since then have been updated based on new computational technologies. Second, the original study included microarray based datasets and not only data derived through RNA-sequencing. Nevertheless, we tried to follow the original methodology as closely as possible, but some of the differences we observed might be attributed to the changes we introduced. In addition, several studies have recently highlighted the issue of the confounding influence of intra-tumoural heterogeneity when using molecular classification[31, 32]. Indeed, a recent study by Li et al, analyzing single cells, has shown that EMT-associated genes, a hallmark of CMS4, are only upregulated in cancer-associated fibroblasts[31]. These findings suggest that CMS4 may not exist as a subtype of CRC per se, rather it may be a reflection of the stromal cells associated with a given tumour. This research area is still under active development and we believe our study provides some important insights into CRC subtyping and associated microbiomes.

Gene-set enrichment analysis showed very similar findings to those described by the CRC subtyping consortium. This is encouraging, as we used a slightly different method to analyse associated biological categories to CRC subtypes (see Materials and Methods). The strong immune signature associated with CMS1, and corresponding low-expression of genes related to immune functions in CMS2 and CMS3, reflect findings of the CRC subtyping consortium[6]. However, a study by Becht et al, found that an inflammatory CRC micro-environmental signature was highly associated with CMS4 in their transcriptomic analysis[33]. In contrast, an inflammatory signature was strongly associated with CMS1 in our study, while CMS2 and CMS3 displayed low immune and inflammatory signals in both studies. Although a small number of studies have used the CMS classifiers on previously unclassified gene array data[34, 35], most published studies reporting of CMS classification have used expression data from the cohorts used in the initial study. This study is the first one to independently validate the classifier using “newly derived” RNA-sequencing data from a physically disparate patient cohort. Our findings highlight the importance of validation of new classification systems in order to ensure their reproducibility and methodological robustness prior to use in a research or clinical setting.

Our second aim involved cataloguing and identifying differences in the tumour microbiome associated with different molecular subtypes of CRC. To this end we employed both 16S rRNA sequencing and RNA-sequencing data. We found significant phylum-level changes between the three subtypes studied in our cohort, using both sequencing methods; differences between the methods may be accounted for by the lower resolution seen using 16S rRNA metabarcoding. Mapping of non-human RNA-sequencing reads to bacterial reference sequences enabled us to catalogue bacterial species of each tumour sample. This identified very distinct bacterial communities associated with each molecular subtype. Although bacterial dysbiosis, as shown through 16S rRNA and metagenomics studies, have previously been associated with CRC compared to controls[9, 10, 36, 37], and a recent study by Burns et al[14] has linked microbial composition with loss-of-function mutations in tumours, this is the first time that different bacterial signatures have been shown to associate with molecular subtypes of CRC.

Increased carriage of *Fusobacterium nucleatum* has been frequently associated with CRC[38-40]. *F. nucleatum* possesses a unique adhesion molecule, FadA, that allows it to adhere to and invade epithelial cells[41]; it has also shown to potentiate colorectal carcinogenesis by recruitment of infiltrating immune cells[42] and modulating E-cadherin/β-catenin signalling[43]. *F. nucleatum* has also been shown to have an association with immune response in the development of CRC[44]. *Fusobacterium hwasookii*, which until recently was classified as *F. nucleatum*[45], was one of the bacterial species most strongly associated with CMS1 in our cohort. Due to its recent reclassification it may account for some of the previously reported associations of *Fusobacterium* with CRC. While the genome of *F. hwasookii* has been sequenced, no mechanistic studies have been carried out to date to compare the oncogenic potential of this strain to *F. nucleatum*. However, given the similarity to the *F. nucleatum* sequence, and presence of a highly conserved *FadA* gene, it is likely that *F. hwasookii* plays a similar role in carcinogenesis. The ability of *Fusobacterium* species to elicit an immune response, in particular to recruit T-cells, is reflected in the immunological signature seen in the CMS1 tumours. Our targeted qPCR analysis of *F. nucleatum* that showed an increased abundance associated with CMS1, also reflects the findings of two studies that found that *Fusobacterium* was associated with a CRC subtype characterised by CpG island methylation, MSI and inflammatory signatures[15], and higher prevalence in right-sided tumours[46], all hallmarks of CMS1[6].

Although not as well studied as *Fusobacterium*, several other oral pathogens, or potential pathobionts, have been reported to be associated with CRC. Of considerable interest to us, is the enrichment of *Porphyromonas gingivalis* with CMS1 in our cohort, identified by both Kraken and qPCR analysis. This oral pathogen has been reported to synergistically promote oral cancer[47] and is associated with CRC, although only some studies reported increased levels of the bacteria in tumours compared to controls[9, 36, 48]. *P. gingivalis* is also a known biofilm former that co-aggregates with *Treponema denticola* and *Tannerrella forsythia*[49, 50], both of which also show enrichment in CMS1. Formation of such biofilms in extra-intestinal infections facilitates synergistic pathogenicity[51, 52], and the high enrichment of these bacteria in CMS1 suggests that similar community synergy may be occurring in the tumour microenvironment. The concept of bacterial biofilms as initiators of CRC has recently been proposed. Biofilms facilitate the invasion of the mucous layer, and a study by Dejea et al[53] found that biofilms were present in > 90% of right-sided CRC; all of the CMS1 tumours in our study were right sided.

We also performed targeted analysis of two further oral bacteria, *P. micra* and *P. stomatis*, in our CRC cohort. These bacterial species have been identified in metagenomics studies as markers of CRC using faecal samples[10], and have been described in an oral-microbe-induced colorectal tumorigenesis model, proposed by Flynn et al[54]. Interestingly, a recent microbiome study by Flemer et al[55] of CRC tumour and matched faecal samples found significantly elevated abundance of *Fusobacterium*, *Peptostreptococcus*, or *Parvimonas* only in a subset of 20-30% of CRC patients. We found enrichment of these bacteria to be associated with CMS1 in our tumour cohort (18%), underlining the potential role of oral polymicrobial communities in the development of a subset of CRC, and the importance of considering CRC heterogeneity when studying mechanisms of CRC pathogenesis.

The major limitation of this study was the small cohort size. Although we found significant associations of bacterial species and taxa associated with particular molecular subgroups, a much larger cohort would be useful to reproduce our findings. We also lacked an independent cohort to carry out validation, thus we carried out validation on the original cohort. Future directions would include testing of qPCR panels in a large independent sample set prior to molecular classification using RNA-sequencing, and investigating the utility of these bacterial markers in non-invasive faecal-based tests.

## Conclusions

In conclusion, due to the potentially modifiable nature of gut bacteria, identifying the role of particular bacterial species in CRC development could have implications for cancer prevention. Here, we have identified, for the first time, distinct microbial populations associated with subtypes of CRC. This will lay the groundwork for future studies into the molecular mechanisms of bacterial colorectal carcinogenesis, and may have clinical utility for CRC screening, diagnosis and treatment.

## Declarations

### Ethics approval and consent to participate

Samples were collected with patients’ written, informed consent, and this study was carried out with approval from the University of Otago Human Ethics Committee (ethics approval number: H16/037).

### Availability of data and material

The datasets used and/or analysed during the current study are available from the corresponding author on reasonable request.

### Funding

#### Funding sources (Rachel Purcell)

Maurice and Phyllis Paykel Trust

Gastrointestinal Cancer Institute (NZ), with support from the Hugh Green Foundation

Colorectal Surgical Society of Australia and New Zealand (CSSANZ)

### Authors' contributions

RP carried out nucleic acid and sequencing preparation of tumour samples and was a major contributor to manuscript writing. MV carried out bioinformatics analysis and preparation of figures for the manuscript. PB was involved in study design and data analysis. SS was involved in study design, bioinformatics analysis and was a major contributor to manuscript preparation. FF was involved in study design and clinical aspects of the study. All authors read and approved the final manuscript.

## Acknowledgements

The authors would like to thank Helen Morrin and the staff at the Cancer Society Tissue Bank, Christchurch for their enthusiasm and support for this project, the patients involved, for generously participating in this study, as well as New Zealand Genomics Limited (NZGL) for the sequencing and support in study design.

